# Machine-learning-guided enzyme discovery and redox-system engineering for efficient production of the nylon monomer methyl 12-aminododecanoate in *Escherichia coli*

**DOI:** 10.64898/2026.09.10.749670

**Authors:** Maiko Umemura, Yasushi Kamisaka, Masaki Yamamoto, Yuki Kuriya, Naoki Watanabe, Chuya Tateishi, Takuya Hashimoto, Manabu Kanno, Shuhei Noda, Michihiro Araki, Kazuhiro Fujimori, Junya Ikuta

## Abstract

Methyl 12-aminododecanoate (ADAME) is a key precursor for Nylon 12 synthesis and an attractive target for sustainable microbial production. However, efficient biosynthesis of ADAME requires the coordinated oxidation and transamination of methyl dodecanoate (DAME), and identifying compatible enzymes for this multistep pathway remains challenging. In this study, we applied a machine learning–based bioinformatic screening approach to identify alternative enzymes for DAME-to-ADAME bioconversion. Among 18 selected distantly related AlkB homologs, AlkBGp01 from *Alcanivorax* sp. P2S70 showed activity toward DAME, producing 12-hydroxydodecanoic acid methyl ester (HDAME) and 12-oxododecanoic acid methyl ester (ODAME) despite low sequence identity to *Pseudomonas putida* AlkB. Further screening identified compatible redox partners, AlkG60 from the same species as AlkBGp01 and AlkTp04, whose combination with AlkBGp01 resulted in almost sixfold higher ODAME production than that achieved with *P. putida* AlkBFGJLT. ODAME production was further increased by more than threefold by conjugating these three newly identified proteins. The optimized single-plasmid system comprising conjugated AlkBGp01, AlkG60, and AlkTp04, together with AlaD and newly identified ω-transaminases EAV41574 produced 0.28 mM gdcw⁻¹ ADAME, representing a 5.6-fold improvement over the previously reported *P. putida* AlkBFGJLT-based system with AlaD–CV2025. These results demonstrate that machine learning–guided enzyme discovery can identify functional distantly related homologs and compatible enzyme combinations for constructing efficient synthetic pathways. This study also highlights the importance of optimizing pathway architecture, including enzyme conjugation and plasmid configuration, for improving microbial production of bio-based monomer precursors.

## 1. Introduction

Nylon 12 (polyamide 12, PA12) is a high-performance engineering plastic widely used in textiles, automotive components, electronics, and medical devices because of its low moisture uptake, dimensional stability, and chemical resistance. However, its industrial production relies almost exclusively on petrochemical feedstocks, motivating the development of sustainable microbial routes to Nylon 12 precursors.

Methyl 12-aminododecanoate (ADAME) is a precursor of the Nylon 12 monomer 12-aminododecanoic acid, representing a promising target for microbial biosynthesis. Previous studies have demonstrated the conversion of methyl dodecanoate (DAME) to ADAME in *Escherichia coli* through heterologous expression of the alkane oxidation system from *Pseudomonas putida*, in combination with ω-transaminase (ω-TA) (Eggink et al., 1987; Schrewe et al., 2013; Ahsan et al., 2018). In this pathway, DAME is converted to ADAME via sequential terminal hydroxylation and oxidation to the corresponding aldehyde catalyzed by the alkane monooxygenase AlkB, rubredoxin AlkG, and rubredoxin reductase AlkT, followed by transamination by ω-TA (Fig. 1). Further improvements in productivity have been achieved by enhancing substrate uptake through the introduction of the outer membrane transporter AlkL (Julsing et al., 2012; van Nuland et al., 2016), as well as by increasing intracellular L-alanine supply via expression of an alanine dehydrogenase (Ladkau et al., 2016).

**Fig. 1.**
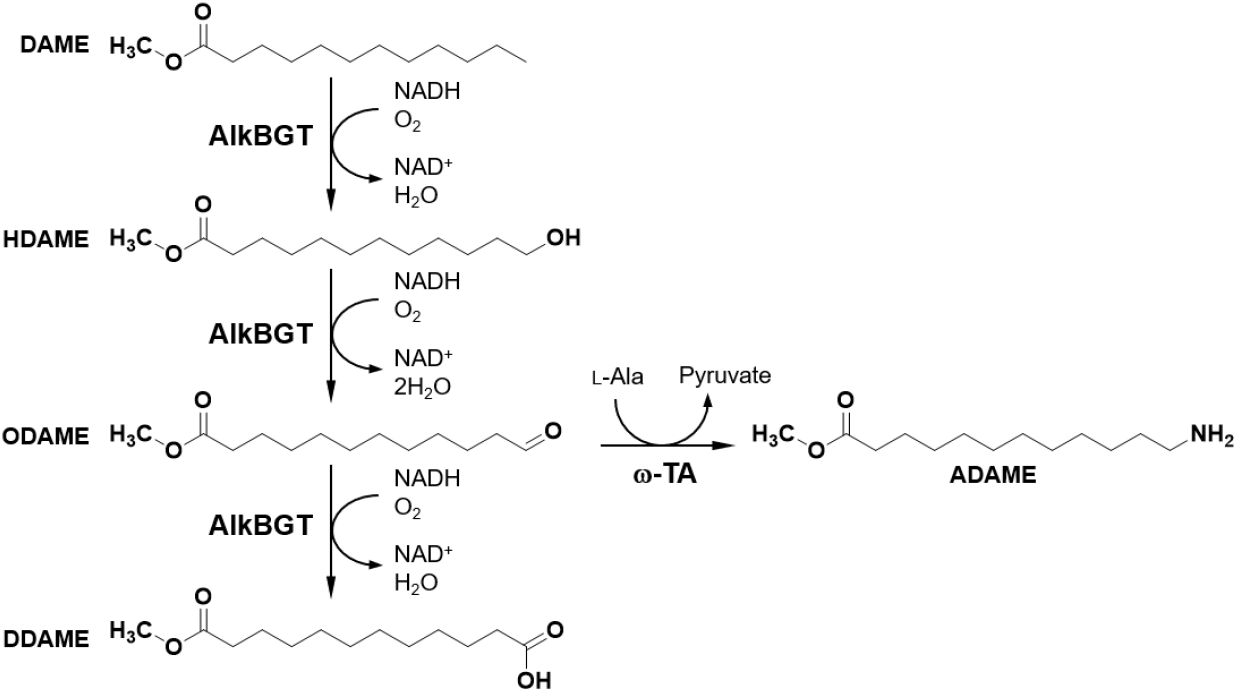
Bioconversion pathway from dodecanoic acid methyl ester (DAME) to 12-aminododecanoic acid methyl ester (ADAME) catalyzed by the AlkBGT alkane oxidation system and ω-transaminase (ω-TA). HDAME, 12-hydroxydodecanoic acid methyl ester; ODAME, 12-oxododecanoic acid methyl ester; DDAME, dodecanedioic acid monomethyl ester.

Despite these advances, the overall efficiency of ADAME production remains limited by several bottlenecks, including restricted substrate uptake, cofactor imbalance, and incomplete oxidation of intermediates. A major limitation is associated with the AlkB-dependent oxidation module, which catalyzes the conversion of DAME to 12-hydroxydodecanoic acid methyl ester (HDAME) and 12-oxododecanoic acid methyl ester (ODAME) (Beilen and Funhoff, 2007). Efficient AlkB catalysis requires coordinated electron transfer from NADH via AlkT and AlkG. Imbalances in protein–protein interactions can therefore reduce electron transfer efficiency, resulting in incomplete oxidation and intermediate accumulation (Ladkau et al., 2016; van Nuland et al., 2016). Furthermore, the membrane-bound nature of AlkB and the soluble characteristics of AlkG and AlkT may limit effective electron transfer, resulting in incomplete oxidation and accumulation of intermediate products.

To address these limitations, we combined machine learning-guided enzyme discovery with pathway engineering to optimize both enzyme compatibility and pathway architecture. We sought to identify compatible homologs of multiple pathway enzymes that collectively improve oxidation efficiency. Alternative homologs of AlkB, AlkT, and ω-TA were identified using a machine learning-based bioinformatic approach. We then engineered compatible AlkBGT complexes by conjugating AlkB with appropriate redox partners to facilitate electron transfer during sequential oxidation. Notably, a naturally occurring fusion enzyme comprising an AlkB-like monooxygenase together with ferredoxin and ferredoxin reductase has been reported (Williams et al., 2022), suggesting that physical coupling of redox partners can enhance electron transfer efficiency. Finally, the optimized oxidation module was integrated with a bioinformatically identified ω-TA and alanine dehydrogenase in a single-plasmid pathway. This work demonstrates the potential of integrating machine learning-guided enzyme discovery with pathway engineering for the development of efficient biosynthetic pathways for bio-based chemicals.

## 2. Materials and methods

### 2.1. Chemicals

DAME (99.8%) and ADAME (99.3%) used as standards were obtained from Fujifilm Wako (Osaka, Japan). HDAME (95%) and ODAME (95%) were obtained from Chemspace (Kiev, Ukraine). All other chemicals used in this work were obtained from Fujifilm Wako (Osaka, Japan) at the Guaranteed Reagent quality unless otherwise specified.

### 2.2. Strains and cultivation

*Escherichia coli* DH5α (Nippon Gene, Tokyo, Japan) was used for cloning and *E. coli* BL21 (DE3) (Nippon Gene, Tokyo, Japan) for biotransformation assays. Recombinant strains were generated by introducing plasmid DNA into chemically competent host cells, and transformants were selected based on antibiotic resistance. Cultivations were performed at 30°C in LCGly1 medium (10 g L⁻¹ LB broth, 18.1 g L⁻¹ Bacto™ CD Supreme Fermentation Production Medium (Thermo Fisher Scientific, Waltham, MA, USA), and 1% glycerol). Where appropriate, antibiotics were added at final concentrations of 100 μg mL⁻¹ sodium carbenicillin and/or 50 μg mL⁻¹ kanamycin sulfate. Solid media contained 1.5% (w/v) agar. Fresh transformants were prepared for each assay.

### 2.3. Construction of enzyme expression vectors

Gene and vector fragments were amplified by PCR using KOD Plus Neo (Toyobo, Osaka, Japan) and Q5 High-Fidelity DNA Polymerase (New England Biolabs, Ipswich, MA, USA), respectively. PCR products were assembled using the NEBuilder HiFi DNA Assembly Master Mix (New England Biolabs) according to the manufacturer’s instructions. Primers were designed to introduce the homologous overlaps required for DNA assembly, including assembly overhangs, native *P. putida* intergenic sequences, or overlapping coding sequences, depending on the construct. For the construction of conjugated genes, internal coding sequences were amplified without their stop codons to generate continuous open reading frames.

The native *P. putida* AlkBFGJLT operon was reconstructed in a pUG6-derived vector (NovoPro Bioscience, Shanghai, China) by assembling four synthetic DNA fragments comprising the *alkBFG* genes with their native intergenic regions and *alkJ*, *alkL*, and *alkT* with their respective native upstream intergenic regions. Deletion constructs lacking *alkB* or *alkT* were generated from this plasmid by inverse PCR using primers flanking the corresponding gene and containing homologous assembly overhangs for subsequent assembly. Synthetic genes encoding the bioinformatically identified enzymes were codon-optimized for *E. coli* and designed with 5′- and 3′-UTRs, containing promoter and terminator sequences and ∼24-bp assembly overhangs for insertion into the corresponding backbone vectors. For plasmid harboring multiple independently expressed *alk* genes, primers were designed to incorporate the corresponding native intergenic regions of the *P. putida alk* operon. Plasmids harboring *alaD*, encoding the alanine dehydrogenase from *Bacillus subtilis* (Ladkau et al., 2016), and bioinformatically identified ω-TA genes were also constructed. Different oxidation modules were subsequently inserted upstream of the *alaD–*ω-TA gene cassette.

Synthetic DNA fragments were synthesized by Eurofins Genomics (Tokyo, Japan). Recombinant plasmids were screened by antibiotic selection and colony PCR and verified by Sanger sequencing of the entire insert region (Eurofins Genomics). Plasmids, *E. coli* strains, and proteins used in this study are listed in Table 1. The sequences of the synthetic genes and primers used in this study are provided in Table S1.

**Table 1.**
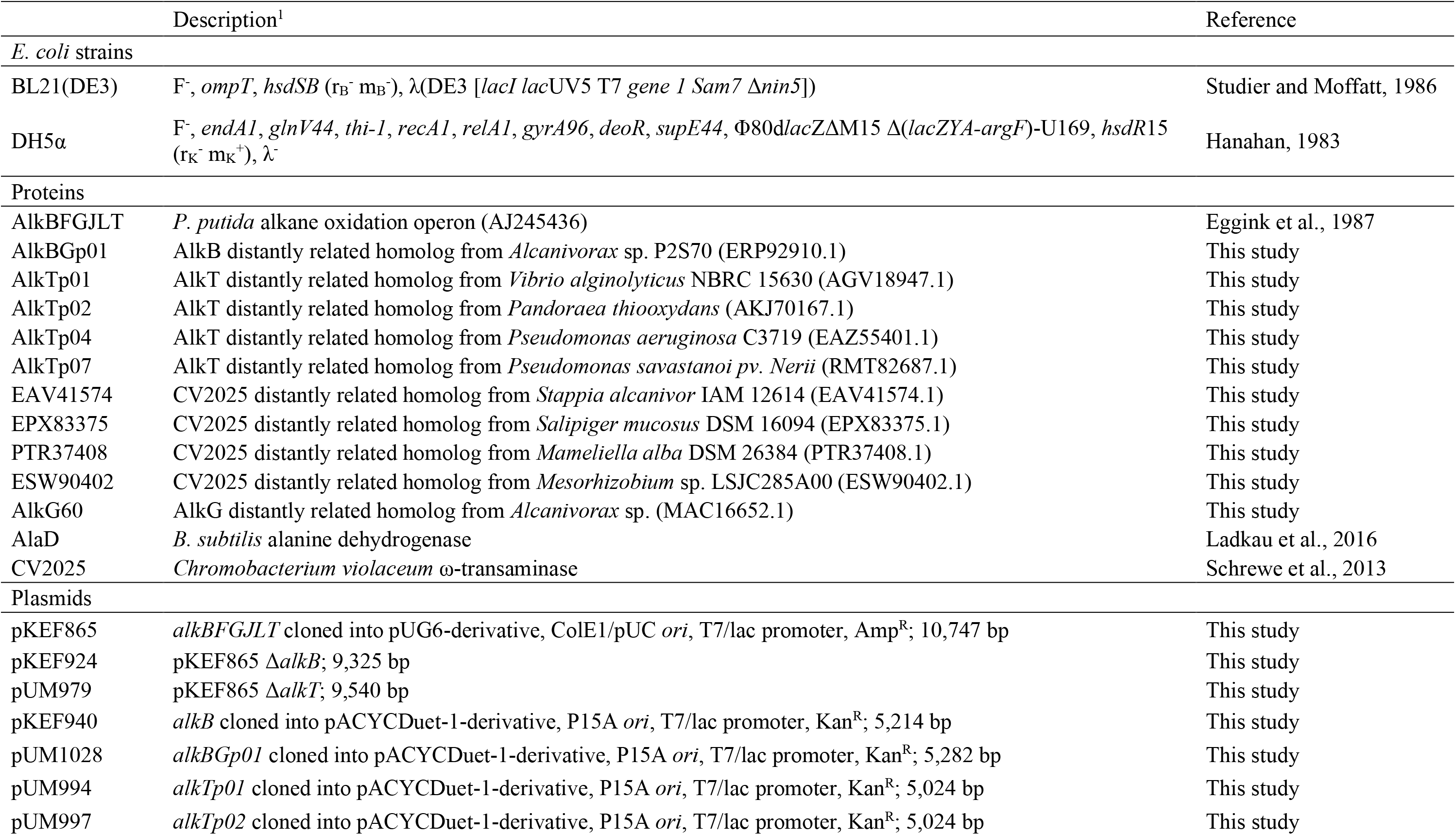
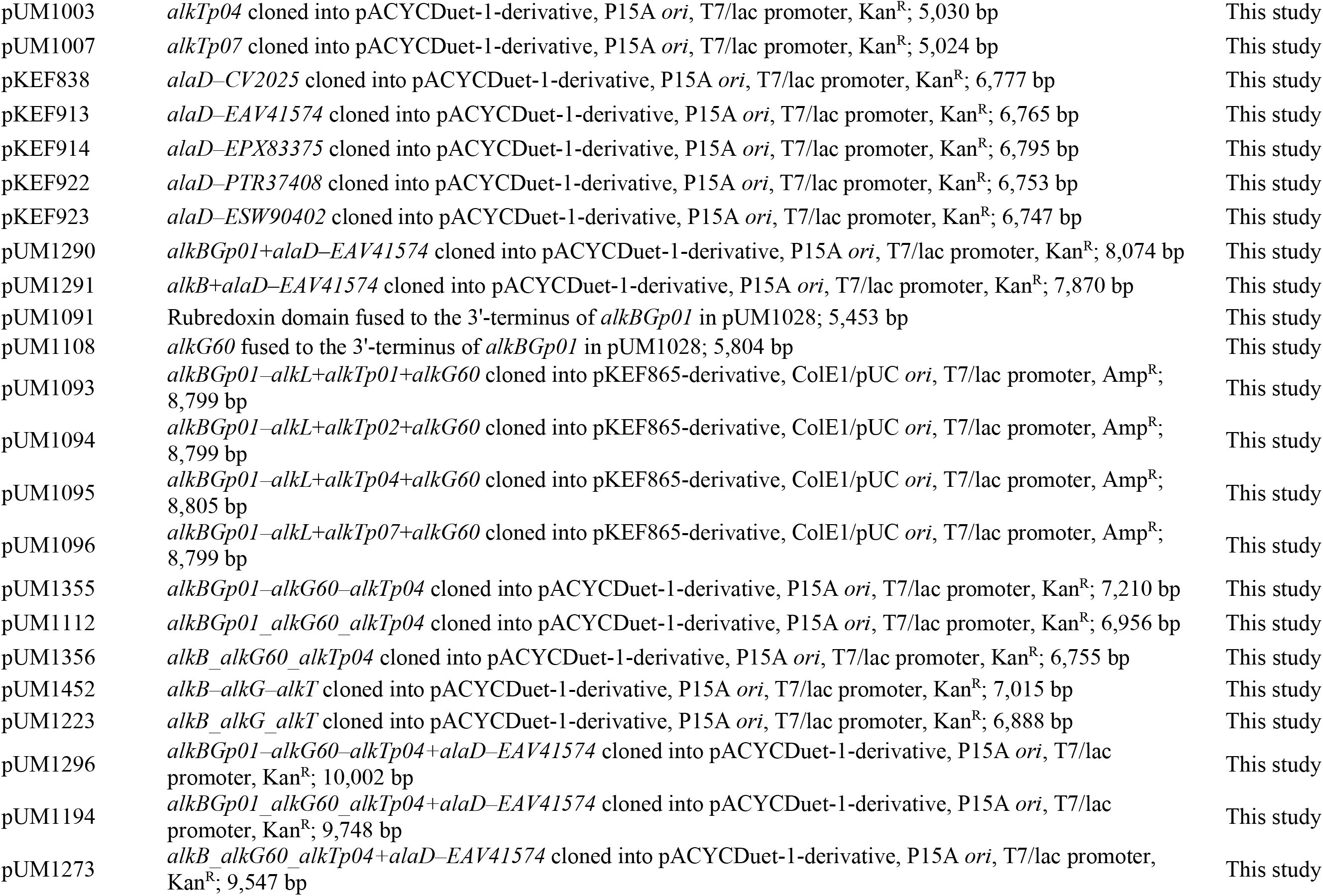
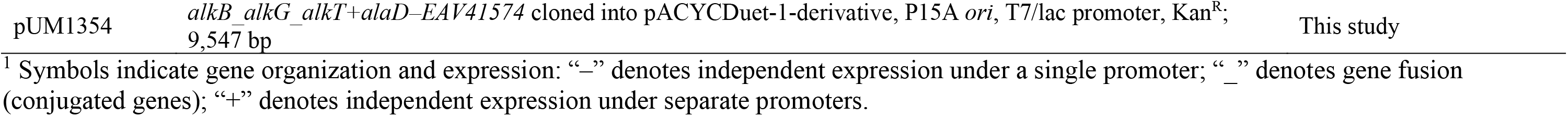
*E. Coli* strains, proteins, and plasmids used in this study.

### 2.4. Biotransformation procedure

For precultivation, 5 mL of LCGly1 medium was inoculated with a single colony from an LB agar plate and incubated for 16 h. Subsequently, 400 μL of the preculture was transferred to 10 mL of LCGly1 medium in a 100 mL baffled flask and incubated with shaking at 200 rpm for 2 h (Sanki Seiki, Osaka, Japan). Protein expression was induced by the addition of 1 mM isopropyl-β-D-thiogalactopyranoside, followed by incubation for 3 h. The cell pellet was harvested by centrifugation at 3,000 rpm for 5 min and resuspended in 0.2 M Tris–HCl buffer (pH 8.0) supplemented with 5% DAME, 2% glucose, 50 mM L-alanine, and 0.1 mM pyridoxal phosphate to obtain a final cell density of 27 gcdw L^-1^. The reaction was carried out in a volume of 200 μL at 30°C with shaking at 2,000 rpm for 2 h. Subsequently, 1 mL of tert-butyl methyl ether containing 0.01% (w/v) methyl eicosanoate (Tokyo Chemical Industry Co., Ltd., Tokyo, Japan) as an internal standard was added, and the mixture was shaken at 2,000 rpm for 10 min. After centrifugation at 15,000 rpm for 30 min, the upper organic phase was collected for gas chromatography–mass spectrometry (GC–MS) analysis of HDAME and ODAME. For derivatizing ADAME prior to GC–MS analysis, 50 μL of N-methyl-bis(trifluoroacetamide) was added to 300 μL of the residual organic phase and heated for 30 min at 60°C. Three biological replicates were performed for each transformant.

### 2.5. Analytic procedures

HDAME, ODAME, and ADAME were analyzed using a GC–MS system (GCMS-QP2010 SE, Shimadzu, Kyoto, Japan) equipped with a DB-FATWAX UI capillary column (30 m × 0.32 mm i.d., 0.25 μm film thickness; Agilent, Santa Clara, CA, USA). A 1 μL sample was injected at 250 °C with a split ratio of 5, using helium as the carrier gas. The oven temperature was initially held at 170 °C for 2 min, then increased to 225 °C at a rate of 2 °C min⁻¹, and held for 1 min, with a total run time of approximately 30 min. The mass spectrometer was operated in electron ionization mode, scanning over an *m*/*z* range of 35–500. The amount of each compound was quantified by comparing peak areas to those of corresponding standards, normalized to the internal standard.

### 2.6. Bioinformatic screening of target enzymes via machine learning

Hypothetical and uncharacterized protein sequences were collected from National Center for Biotechnology Information (NCBI) non-redundant (nr) protein database as of 2019 (Sayers *et al*., 2019). The sequences that were duplicated and included non-canonical amino acids were removed. The search data for each enzyme candidate is shown as Table 2. To ensure the quality of the candidates, the range of amino acid counts was independently determined for each enzyme type.

**Table 2.** Dataset information of bioinformatic screening.

| Enzyme | EC number | Sequence length | Number of prediction samples | Number of samples in first screening | Number of final candidates |
| --- | --- | --- | --- | --- | --- |
| AlkB | EC 1.14.15.X | 250-550aa | 11.9 M | 295 | 75 |
| AlkT | EC 1.18.1.1 | 350-500aa | 2.37 M | 330 | 10 |
| ω-TA | EC 2.6.1.X | 350-550aa | 0.47 M | 913 | 10 |

\*M: Million; All sequences were retrieved from the NCBI nr database as of 2019.

For each dataset, Enzyme Commission numbers (ECs) were predicted using EnzymeNet (Watanabe *et al*., 2023). The tool first identified the primary EC class and then assigned the complete four-digit EC from protein sequences. From the outputs, sequences that matched the target enzyme’s EC at both the first- and fourth-digit levels with prediction scores of 0.5 or higher for each step were extracted as the primary screening for enzyme candidates.

Following the initial screening, a four-digit EC model was used to encode both the candidate sequences and the reference enzyme sequences into 2,048-dimensional feature vectors. The similarity between the feature vectors was quantified using the Tanimoto similarity score, with higher scores indicating greater similarity in the learned feature space rather than amino acid sequence identity. Sequences with high Tanimoto similarity scores to the reference enzyme vectors were prioritized as potential candidates. For AlkB, experimentally validated inactive analogs were also included as reference sequences. In addition to the similarity-based criteria, candidates for AlkB were filtered by comparing their feature vectors with the centroids of the active and inactive groups; only sequences positioned closer to the active enzyme centroid than to the inactive centroid were selected.

## 3. Results

### 3.1. Activity screening of bioinformatically selected AlkB, AlkT, and ω-transaminase homologs

Bioinformatic screening using our machine learning-based approach identified 75 proteins remotely homologous to *P. putida* AlkB with Tanimoto similarity scores above 0.557. From these, 18 top-ranked candidates were selected for biotransformation assays after excluding highly similar sequences. The selected genes were cloned into expression vectors and coexpressed in *E. coli* BL21(DE3) with a second plasmid harboring the *P. putida* alkane oxidation operon components AlkFJGLT. AlkF is an additional rubredoxin, and AlkJ is an alcohol dehydrogenase. Among the candidates, only one—ERP92910.1 from *Alcanivorax* sp. P2S70 (AlkBGp01)—exhibited biotransformation activity from DAME to HDAME and ODAME. AlkBGp01 produced less than one-third of the HDAME and approximately 80% of the ODAME levels compared with *P. putida* AlkB coexpressed with the AlkFJGLT plasmid (Table 3). AlkBGp01 shares 47% sequence identity with *P. putida* AlkB.

**Table 3.** Biotransformation yields of bioinformatically identified enzymes.

| Construct | Protein(s) | Yield (mM g<sub>DCW</sub><sup>-1</sup>) |  |  |  |
| --- | --- | --- | --- | --- | --- |
|  |  | HDAME | ODAME | ADAME | Total |
| pKEF865 | AlkBFGJLT | 0.57 ± 0.05 | 0.10 ± 0.01 | - | 0.67 |
| pKEF924/pKEF940 | AlkFJGLT/AlkB | 1.96 ± 0.08 | 0.50 ± 0.02 | - | 2.46 |
| pKEF924/pUM1028 | AlkFJGLT/AlkBGP01 | 0.73 ± 0.09 | 0.41 ± 0.02 | - | 1.14 |
| pUM979/pUM982 | AlkBFGJL/AlkT | 0.82 ± 0.05 | 0.10 ± 0.01 | - | 0.92 |
| pUM979/pUM994 | AlkBFGJL/AlkTp01 | 1.55 ± 0.01 | 0.45 ± 0.01 | - | 2.00 |
| pUM979/pUM997 | AlkBFGJL/AlkTp02 | 1.93 ± 0.09 | 0.64 ± 0.04 | - | 2.57 |
| pUM979/pUM1003 | AlkBFGJL/AlkTp04 | 1.62 ± 0.13 | 0.61 ± 0.03 | - | 2.23 |
| pUM979/pUM1007 | AlkBFGJL/AlkTp07 | 1.36 ± 0.03 | 0.31 ± 0.01 | - | 1.67 |
| pKEF865/pKEF838 | AlkBFGJLT/AlaD-CV2025 | 1.08 ± 0.04 | 0.11 ± 0.02 | 0.05 ± 0.01 | 1.24 |
| pKEF865/pKEF913 | AlkBFGJLT/AlaD-EAV41574 | 0.91 ± 0.02 | 0.06 ± 0.01 | 0.06 ± 0.01 | 1.03 |
| pKEF865/pKEF914 | AlkBFGJLT/AlaD-EPX83375 | 1.03 ± 0.05 | 0.12 ± 0.01 | 0.06 ± 0.00 | 1.21 |
| pKEF865/pKEF922 | AlkBFGJLT/AlaD-PTR37408 | 1.07 ± 0.04 | 0.14 ± 0.01 | 0.09 ± 0.00 | 1.30 |
| pKEF865/pKEF923 | AlkBFGJLT/AlaD-ESW90402 | 1.08 ± 0.05 | 0.12 ± 0.02 | 0.06 ± 0.00 | 1.26 |

<sup>1</sup> Symbols indicate gene organization and expression: “—” denotes independent expression under a single promoter; “/” denotes expression from separate plasmids.

Using the same bioinformatic screening approach as for AlkB, 10 remotely homologous AlkT proteins were identified with Tanimoto similarity scores above 0.811, of which 8 were selected for biotransformation assays after excluding highly similar sequences. These genes were cloned into expression vectors and coexpressed with a second plasmid harboring *P. putida* AlkBFGJL. Four candidates—AlkTp01, AlkTp02, AlkTp04, and AlkTp07—exhibited 1.8- to 2.8-fold higher total production of HDAME and ODAME than *P. putida* AlkT (Table 3). These proteins were derived from *Vibrio alginolyticus*, *Pandoraea thiooxydans*, *Pseudomonas aeruginosa*, and *Pseudomonas savastanoi*, with sequence identities of 38%, 39%, 55%, and 57% to *P. putida* AlkT, respectively.

For ω-TA screening, 13 proteins with Tanimoto similarity scores above 0.808 were identified, and 10 candidates were selected for biotransformation assay. The selected genes were cloned into expression vectors downstream of *alaD* and coexpressed with another plasmid harboring *P. putida* AlkBFGJLT (pKEF865). Four candidates—EAV41574, EPX83375, PTR37408, and ESW90402—showed more ADAME production than the *Chromobacterium violaceum* ω-TA CV2025 used in the previous study (Schrewe et al., 2013) (Table 3). These proteins originated from *Stappia aggregata*, *Salipiger mucosus*, *Mameliella alba*, and *Mesorhizobium* sp., with sequence identities of 33%, 54%, 36%, and 33% to CV2025, respectively.

In all cases, the bioinformatically identified enzymes exhibited low sequence identity and originated from strains phylogenetically distant from *P. putida* or *C. violaceum*, suggesting that they would be difficult to select as candidates using conventional homology searches. These results demonstrate that the machine learning-based approach effectively identifies distantly related homologs with the desired enzymatic activity.

### 3.2. Bioconversion activity of conjugated enzymes identified through bioinformatics

Among the 18 bioinformatically identified alternative AlkB candidates, the AlkB distantly related homolog from *Alcanivorax* sp. P2S70 (AlkBGp01), which showed the highest Tanimoto score, was the only enzyme exhibiting DAME biotransformation activity. We first compared the conversion efficiency from DAME to ADAME between *E. coli* systems expressing either AlkBGp01 or AlkB, together with AlaD and one of the bioinformatically identified ω-TA, EAV41574, and coexpressed with a second plasmid harboring *P. putida* AlkFGJLT. The system containing AlkBGp01 showed 1.2-fold higher levels of ADAME than that containing AlkB, although the levels of intermediate products (HDAME and ODAME) were lower (Table 4). These results suggest that AlkBGp01 is a promising alternative to AlkB for more efficient ADAME production in *E. coli*.

**Table 4.** Biotransformation yields of AlkBGp01 in combination with bioinformatically identified partner enzymes.

| Construct | Protein(s) | Yield (mM<br>g<sub>DCW</sub><sup>-1</sup>) |  |  |  |
| --- | --- | --- | --- | --- | --- |
|  |  | HDAME | ODAME | ADAME | Total |
| pKEF924/pUM1291 | AlkFGJLT/AlkB+AlaD-EAV41574 | 1.17 ± 0.02 | 0.15 ± 0.02 | 0.23 ± 0.02 | 1.55 |
| pKEF924/pUM1290 | AlkFGJLT/AlkB<sub>Gp01</sub>+AlaD-EAV41574 | 0.87 ± 0.06 | 0.09 ± 0.03 | 0.27 ± 0.01 | 1.23 |
| pKEF924/pUM1091 | AlkFGJLT/AlkB<sub>Gp01</sub>_gp01 | 0.72 ± 0.05 | 0.23 ± 0.01 | - | 0.95 |
| pKEF924/pUM1108 | AlkFGJLT/AlkB<sub>Gp01</sub>_AlkG60 | 1.29 ± 0.06 | 0.69 ± 0.03 | - | 1.98 |
| pUM1093/pKEF913 | AlkB<sub>Gp01</sub>-AlkL+AlkTp01+AlkG60/AlaD-EAV41574 | 0.91 ± 0.04 | 0.15 ± 0.01 | 0.02 ± 0.00 | 1.07 |
| pUM1094/pKEF913 | AlkB<sub>Gp01</sub>-AlkL+AlkTp02+AlkG60/AlaD-EAV41574 | 0.55 ± 0.03 | 0.11 ± 0.02 | 0.00 ± 0.00 | 0.66 |
| pUM1095/pKEF913 | AlkB<sub>Gp01</sub>-AlkL+AlkTp04+AlkG60/AlaD-EAV41574 | 0.61 ± 0.05 | 0.06 ± 0.00 | 0.11 ± 0.01 | 0.78 |
| pUM1096/pKEF913 | AlkB<sub>Gp01</sub>-AlkL+AlkTp07+AlkG60/AlaD-EAV41574 | 0.55 ± 0.06 | 0.02 ± 0.00 | 0.09 ± 0.01 | 0.67 |
| pUM1355 | AlkB<sub>Gp01</sub>-AlkG60-AlkTp04 | 0.53 ± 0.27 | 0.18 ± 0.07 | - | 0.71 |
| pUM1112 | AlkB<sub>Gp01</sub>_AlkG60_AlkTp04 | 0.64 ± 0.05 | 0.58 ± 0.01 | - | 1.23 |
| pUM1452 | AlkB-AlkG-AlkT | 0.31 ± 0.02 | 0.02 ± 0.01 | - | 0.34 |
| pUM1223 | AlkB_AlkG_AlkT | 0.06 ± 0.02 | 0.00 ± 0.00 | - | 0.06 |
| pUM1356 | AlkB_AlkG60_AlkTp04 | 0.29 ± 0.02 | 0.07 ± 0.02 | - | 0.36 |
| pUM1296 | AlkB<sub>Gp01</sub>-AlkG60-AlkTp04+AlaD-EAV41574 | 0.60 ± 0.04 | 0.04 ± 0.01 | 0.09 ± 0.01 | 0.73 |
| pUM1194 | AlkB<sub>Gp01</sub>_AlkG60_AlkTp04+AlaD-EAV41574 | 0.83 ± 0.04 | 0.36 ± 0.02 | 0.28 ± 0.02 | 1.48 |
| pUM1273 | AlkB_AlkG60_AlkTp04+AlaD-EAV41574 | 0.40 ± 0.02 | 0.05 ± 0.02 | 0.05 ± 0.00 | 0.50 |
| pUM1354 | AlkB_AlkG_AlkT+AlaD-EAV41574 | 0.06 ± 0.01 | 0.00 ± 0.00 | 0.00 ± 0.00 | 0.06 |

<sup>1</sup> Symbols indicate gene organization and expression: “-” denotes independent expression under a single promoter; “\_” denotes gene fusion
(conjugated genes); “+” denotes independent expression under separate promoters; “/” denotes expression from separate plasmids.

AlkBGp01 contains a rubredoxin domain at its C-terminus (Fig. 2a). Notably, *P. putida* AlkG contains two tandem rubredoxin domains, AlkG1 and AlkG2, connected by a spacer sequence; AlkG2 is capable of electron transfer, whereas AlkG1 is not, due to an arginine insertion immediately following the second catalytic CXXCX motif (van Beilen et al., 2002). Sequence analysis indicated that the native C-terminal rubredoxin domain of AlkBGp01 belongs to the AlkG1 type (Fig. S1a), suggesting that it is unlikely to function directly in electron transfer but may nevertheless contribute to the oxidation activity of AlkBGp01. To examine whether an additional copy of this domain could enhance AlkBGp01 activity, we fused a second copy of its native C-terminal rubredoxin domain to the C-terminus of AlkBGp01 (AlkBGp01_gp01). However, the resulting construct did not improve bioconversion of DAME to HDAME and ODAME compared with AlkBGp01 alone. The HDAME yield was comparable between AlkBGp01 and AlkBGp01_gp01, whereas ODAME production decreased by approximately 44%, from 0.41 to 0.23 mM gdcw^-1^ (Table 3 and Table 4).

**Fig. 2.**
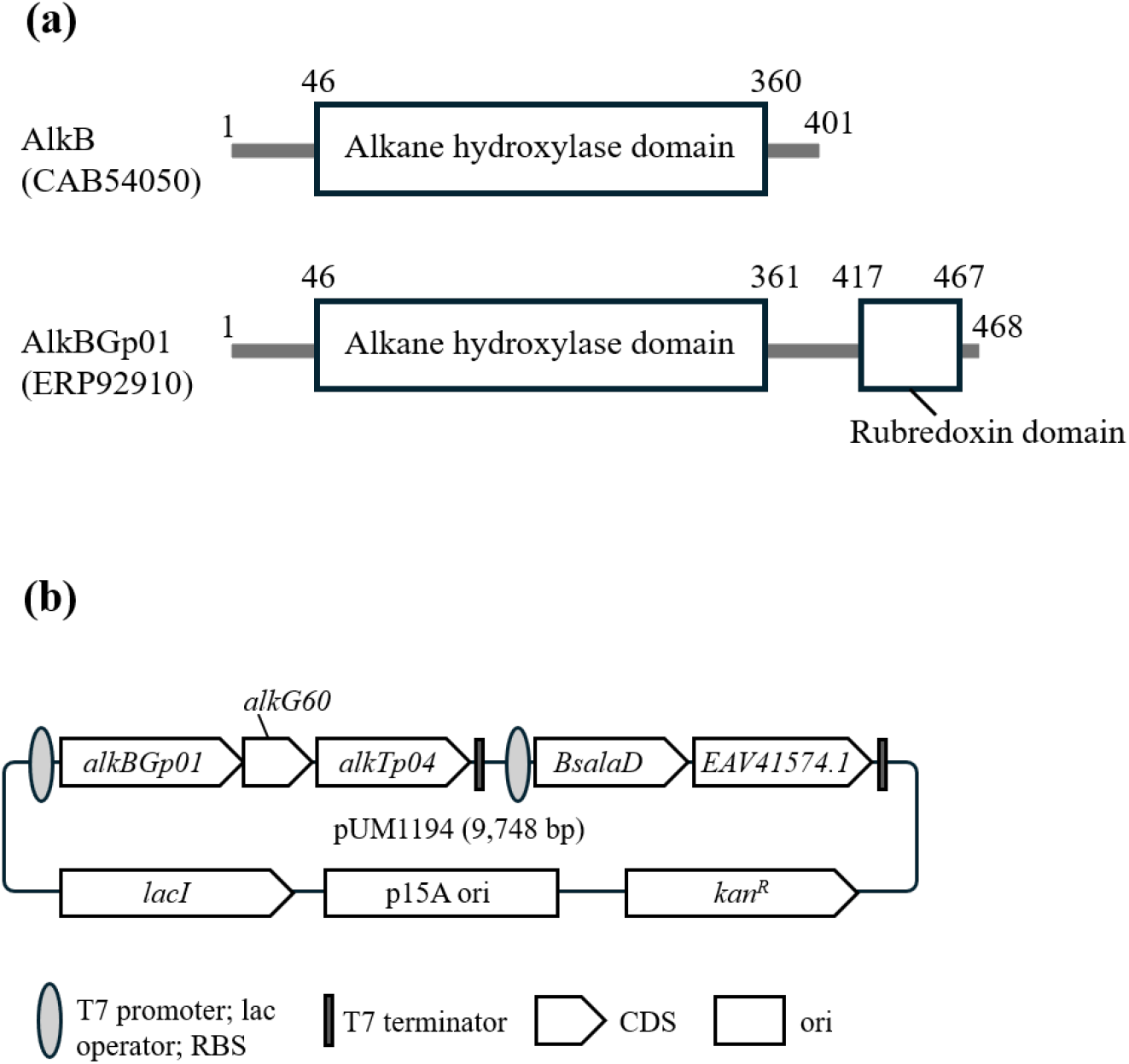
Functional domain architectures of *Pseudomonas putida* AlkB and *Alcanivorax* sp. P2S70 AlkBGp01 (A), and the architecture of the final plasmid pUM1194 for bioconversion of dodecanoic acid methyl ester (DAME) to 12-aminododecanoic acid methyl ester (ADAME) (B).

We then hypothesized that rubredoxin from the same species as AlkBGp01 might provide better compatibility with AlkGBp01. Accordingly, an AlkG homolog from *Alcanivorax* sp. (MAC16652.1; AlkG60), which shares 55.37% sequence identity with *P. putida* AlkG, was fused to the C-terminus of AlkBGp01 (AlkBGp01_AlkG60) and coexpressed with another plasmid harboring AlkFGJLT. This fusion construct increased the production of both HDAME and ODAME, resulting in a 1.7-fold increase in total product yield compared with AlkBGp01 alone (Table 3 and Table 4). These results suggest that coordinated coupling of AlkBGp01 with the compatible redox partner enhanced catalytic efficiency in the alkane oxidation cascade.

We next evaluated four bioinformatically identified AlkT homologs—AlkTp01, AlkTp02, AlkTp04, and AlkTp07—for compatibility with AlkBGp01. Before constructing conjugated enzymes, each AlkT homolog was tested in an individually expressed system together with AlkBGp01 and AlkG60 for the bioconversion of DAME to HDAME, ODAME, and ADAME. To ensure sufficient expression, AlkBGp01, the AlkT homolog, and AlkG60 were each expressed under separate promoters and coexpressed with a second plasmid harboring *alaD* and *EAV41574*. To simplify the construct, AlkF, AlkJ, and AlkL were omitted because they did not consistently improve ADAME production under the conditions tested. Among the four homologs, AlkTp01 and AlkTp02 exhibited negligible ADAME production, whereas AlkTp04 and AlkTp07 produced appreciable levels of ADAME (0.11 and 0.09 mM g_DCW_^-1^, respectively) (Table 4). AlkTp04, showing the highest ADAME production, was therefore selected for subsequent conjugation with AlkBGp01 and AlkG60 in the order of electron transfer.

In an *E. coli* system lacking transamination enzymes, the conjugated AlkBGp01_AlkG60_AlkTp04 construct produced 1.2-fold more HDAME (0.64 mM g_DCW_^-1^) and 3.2-fold more ODAME (0.58 mM g_DCW_^-1^) than the corresponding independently expressed system (Table 4). In contrast, conjugation of *P. putida* AlkB, AlkG, and AlkT markedly reduced HDAME production from 0.31 to 0.06 mM g_DCW_^-1^ and completely abolished ODAME production (Table 4). Similarly, when *P. putida* AlkB was conjugated with AlkG60 and AlkTp04, HDAME production decreased to less than half and ODAME production to approximately one-eighth of the levels observed with the conjugated AlkBGp01_AlkG60_AlkTp04 system (Table 4). These results indicate that AlkBGp01, AlkG60, and AlkTp04 constitute a compatible electron-transfer system that promotes oxidation beyond the HDAME intermediate to ODAME.

We further evaluated the compatibility of these constructs with the bioinformatically identified ω-TA EAV41574 for ADAME production by constructing single-plasmid systems in which the transamination module, AlaD–EAV41574, was cloned downstream of each alkane oxidation module. Consistent with the above results, the conjugated AlkBGp01_AlkG60_AlkTp04 system containing the transamination module produced 3.1-fold more ADAME than the corresponding independently expressed system (0.28 vs. 0.09 mM g_DCW_^-1^) (Table 4). Moreover, the HDAME-to-ODAME ratio was approximately 2:1 in the conjugated system, compared with 15:1 in the independently expressed system, indicating that oxidation proceeded efficiently to ODAME rather than stalling at the HDAME intermediate. The conjugated system containing *P. putida* AlkB (AlkB_AlkG60_AlkTp04) also showed substantial HDAME accumulation, with an HDAME-to-ODAME ratio of 8:1, resulting in markedly reduced ADAME production (0.05 mM g_DCW_^-1^). These results suggest that efficient oxidation to ODAME depends not solely on AlkG60, but on its compatibility with AlkBGp01. The conjugated *P. putida* AlkB_AlkG_AlkT containing the transamination module produced low levels of HDAME and no detectable ODAME or ADAME.

Collectively, these results demonstrate that the bioinformatically identified AlkBGp01, when conjugated with appropriate redox partner proteins, enables more efficient conversion of DAME to ADAME than the native *P. putida* AlkBGT system. Figure 3 summarizes the biotransformation yields of HDAME, ODAME, and ADAME obtained with representative oxidation and transamination systems. The system containing *P. putida* AlkBFGJLT with AlaD and CV2025 showed inefficient conversion of HDAME to ADAME, as reflected by an HDAME-to-ADAME ratio of 22:1. In contrast, the conjugated AlkBGp01_AlkG60_AlkTp04 system, together with AlaD and the bioinformatically identified ω-TA EAV41574, increased ADAME production 5.6-fold compared with the previously reported system (0.28 vs 0.05 mM g_DCW_^-1^) (Fig. 3; Table 3 and Table 4). The architecture of the final single plasmid, pUM1194, is shown in Fig. 2b. This plasmid is 9,748 bp and contains alkane oxidation and transamination modules, each individually controlled by a T7 promoter.

**Fig. 3.**
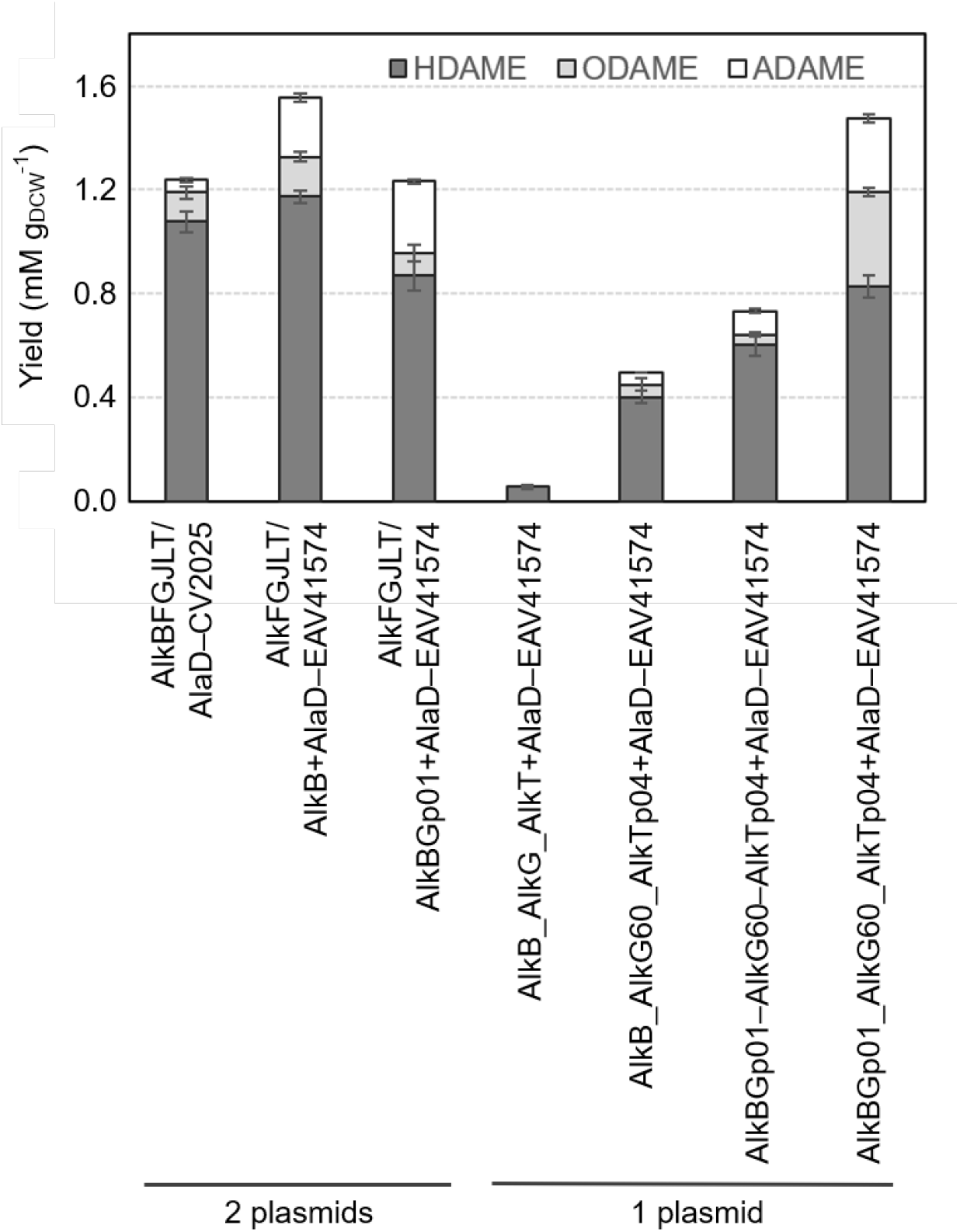
Biotransformation yields of HDAME, ODAME, and ADAME obtained with representative alkane oxidation–transamination systems. Symbols in the construct names indicate gene organization and expression: “–” denotes independent expression under a single promoter; “_” denotes gene fusion; “+” denotes independent expression under separate promoters; and "/" denotes expression from separate plasmids.

## 4. Discussion

The present study demonstrates that combining machine learning-guided enzyme discovery with pathway engineering can substantially improve ADAME production. It also highlights that successful engineering of the DAME-to-ADAME pathway requires not only the identification of suitable enzymes but also optimization of their compatibility within the pathway.

Among the 18 alternative AlkB candidates tested, only one enzyme, AlkBGp01, exhibited alkane oxidation activity. AlkB and AlkBGp01 were both predicted to contain six transmembrane helices by DeepTMHMM (Hallgren et al., 2022), consistent with the crystal structures of FtAlkB and FtAlkBG from *Fontimonas thermophila* (PDB Ids: 8SBB and 8F6T, respectively) (Chai et al., 2023; Guo et al., 2023). Besides the additional C-terminal rubredoxin domain and their low overall sequence identity, the major structural difference between AlkBGp01 and AlkB is a four-amino-acid insertion between transmembrane helices TM1 and TM2 (Fig. S1b). Structural comparison with FtAlkBG suggests that this insertion (LFEQ) is located on the periplasmic side of the membrane (Fig. 4a), whereas the fused rubredoxin domain is on the cytoplasmic side. Therefore, the insertion is unlikely to directly interfere with electron transfer between AlkBGp01 and its fused redox partner, although its functional significance remains unclear.

**Fig. 4.**
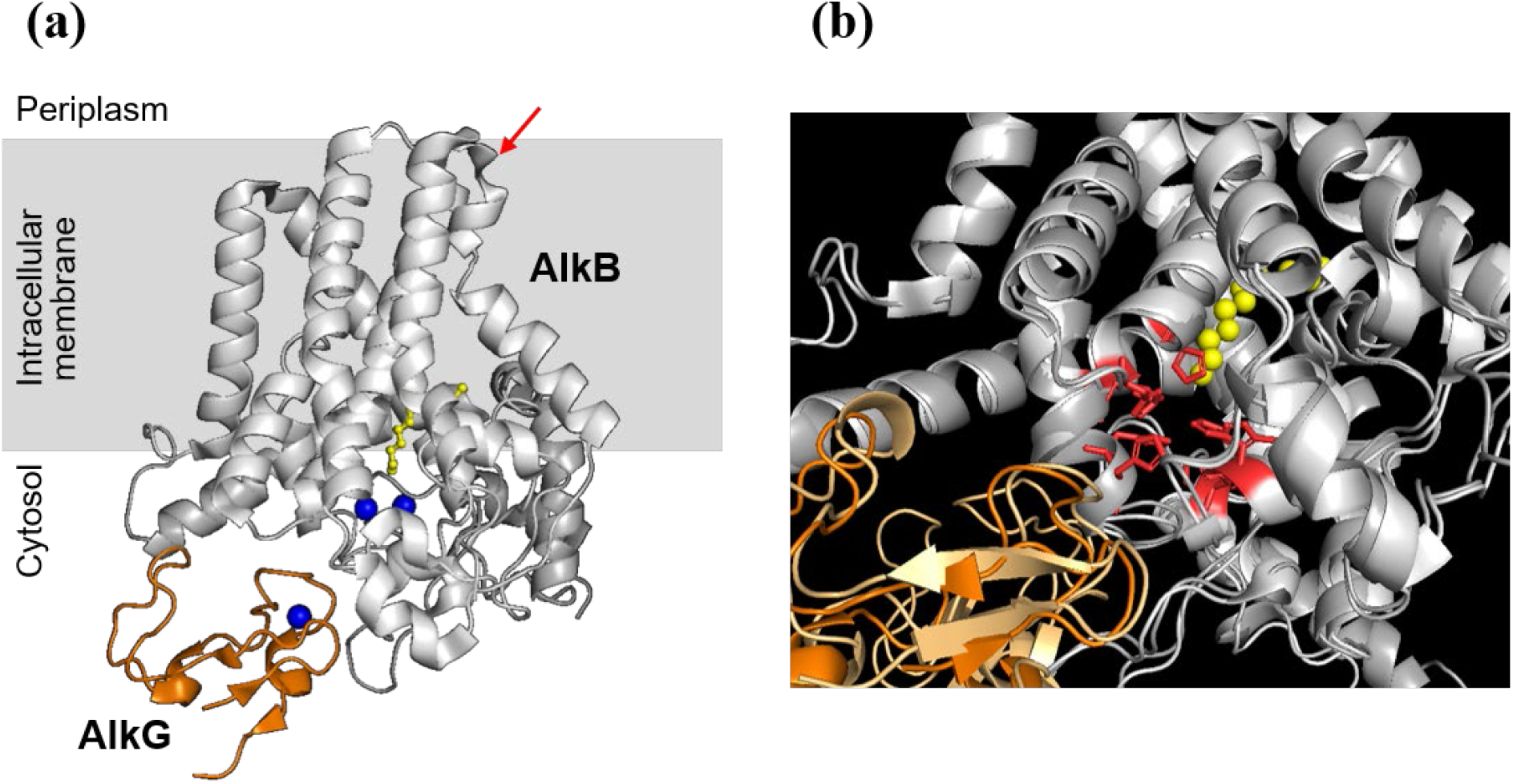
Structures of the AlkB–G complex. (a) Structure of FtAlkBG (PDBID: 8F6T). The red arrow indicates the position corresponding to the four-residue insertion in AlkBGp01. The ligand is shown in yellow and the three Fe ions in blue. (b) Superposition of the catalytic site of FtAlkBG with that of the AlkBGp01–AlkG60 complex predicted by AlphaFold3. FtAlkB and AlkBGp01 are shown in light and dark grey, respectively, whereas FtAlkG and AlkG60 are shown in orange and light orange, respectively. The nine His residues in the AlkB catalytic center are shown in red.

AlkG60 shares only 55.37% sequence identity with *P. putida* AlkG, but most of the sequence divergence is located within the spacer region separating the two rubredoxin domains (van Beilen et al., 2002). The electron-transfer-competent Type G2 rubredoxin domain is 92% conserved between AlkG60 and AlkG (Fig. S1a). Nevertheless, replacing AlkBGp01 with *P. putida* AlkB in the conjugated AlkBGp01_AlkG60_AlkTp04 architecture markedly reduced oxidation efficiency, particularly ODAME and ADAME production (Fig. 3; Table 4). This result suggests that AlkBGp01 is structurally more compatible with AlkG60 than *P. putida* AlkB is, particularly around the catalytic site. In the AlkB−AlkG complex, the alkane oxidation catalytic site of AlkB is located adjacent to AlkG, with nine His residues in AlkB coordinating two Fe ions and four Cys residues in AlkG coordinating one Fe ion. The predicted structure of the AlkBGp01−AlkG60 complex by AlphaFold3 (Abramson et al., 2024) suggests that AlkG60 is positioned closer to the catalytic site of AlkBGp01 than the corresponding *P. putida* AlkB−AlkG complex (Fig. 4b).

Although machine learning enabled the discovery of distantly related homologs with the desired catalytic activities, overall productivity depended strongly on the genetic architecture of the expression system. For example, the *P. putida* AlkBFGJLT system together with AlaD– CV2025 exhibited substantially higher ADAME production when *alkB* was removed from the *alk* operon and expressed from a separate plasmid together with the transamination module (Fig. 3; Table 4). In both systems, *alkB* was positioned immediately downstream of the T7 promoter, whereas the plasmids harboring *alkB* were high- and medium-copy-number vectors, respectively. This observation suggests that either maximal expression of *alkB* or the expression balance among AlkB, AlkG, and AlkT may not be optimal in the former system, possibly because AlkB is an inner membrane protein. Similar effects were observed for multiple pathway configurations, highlighting the strong influence of genetic architecture on pathway performance. However, comprehensive optimization was impractical because the pathway comprises five configurable enzymes, resulting in a vast number of possible combinations of gene order, promoter organization, plasmid copy number, and enzyme conjugation. These results suggest that, in addition to enzyme discovery, systematic optimization of pathway architecture represents an important opportunity for future machine learning-guided metabolic engineering.

## 5. Conclusion

In this study, we demonstrated the utility of our machine learning-based approach for identifying alternative enzymes for the bioconversion of DAME to ADAME, a key precursor for Nylon 12 synthesis. The newly identified enzymes, AlkBGp01, AlkTp04, and ω-TA EAV41574, share low sequence identity with their previously reported counterparts, demonstrating the ability of our approach to identify functional distantly related homologs. By combining these enzymes with AlkG60 and alanine dehydrogenase in a single engineered pathway, we achieved substantially improved ADAME production compared with the previously reported *P. putida* AlkBGT-based system. These results highlight the potential of machine learning-guided enzyme discovery for constructing efficient synthetic metabolic pathways.

## Acknowledgements

This research was based on results obtained from a project, JPNP20011, commissioned by the New Energy and Industrial Technology Development Organization (NEDO).

## CRediT authorship contribution statement

Maiko Umemura: Conceptualization, Methodology, Formal analysis, Supervision, Writing – original draft, Writing – review & editing.

Yasushi Kamisaka: Methodology, Investigation, Formal analysis, Writing – original draft, Writing – review & editing.

Masaki Yamamoto: Methodology, Formal analysis, Software.

Yuki Kuriya: Methodology, Formal analysis, Software. Naoki Watanabe: Methodology, Formal analysis.

Chuya Tateishi: Formal analysis, Investigation, Writing – review & editing. Takuya Hashimoto: Formal analysis, Writing – review & editing.

Manabu Kanno: Formal analysis, Writing – review & editing. Shuhei Noda: Formal analysis.

Michihiro Araki: Conceptualization, Methodology, Supervision, Formal analysis, Software, Writing – review & editing.

Kazuhiro Fujimori: Conceptualization, Supervision, Methodology, Investigation, Formal analysis.

Junya Ikuta: Project administration, Conceptualization, Supervision, Methodology, Investigation, Formal analysis, Writing – review & editing.

## Data availability

Data supporting the findings of this study are available within the article and its Supplementary Material.

## Abbreviations

ADAME: methyl 12-aminododecanoate
DAME: methyl dodecanoate
HDAME: 12-hydroxydodecanoic acid methyl ester
ODAME: 12-oxododecanoic acid methyl ester
ω-TA: ω-transaminase
GC–MS: gas chromatography–mass spectrometry
EC: Enzyme Commission number

---

**Fig. S1.**
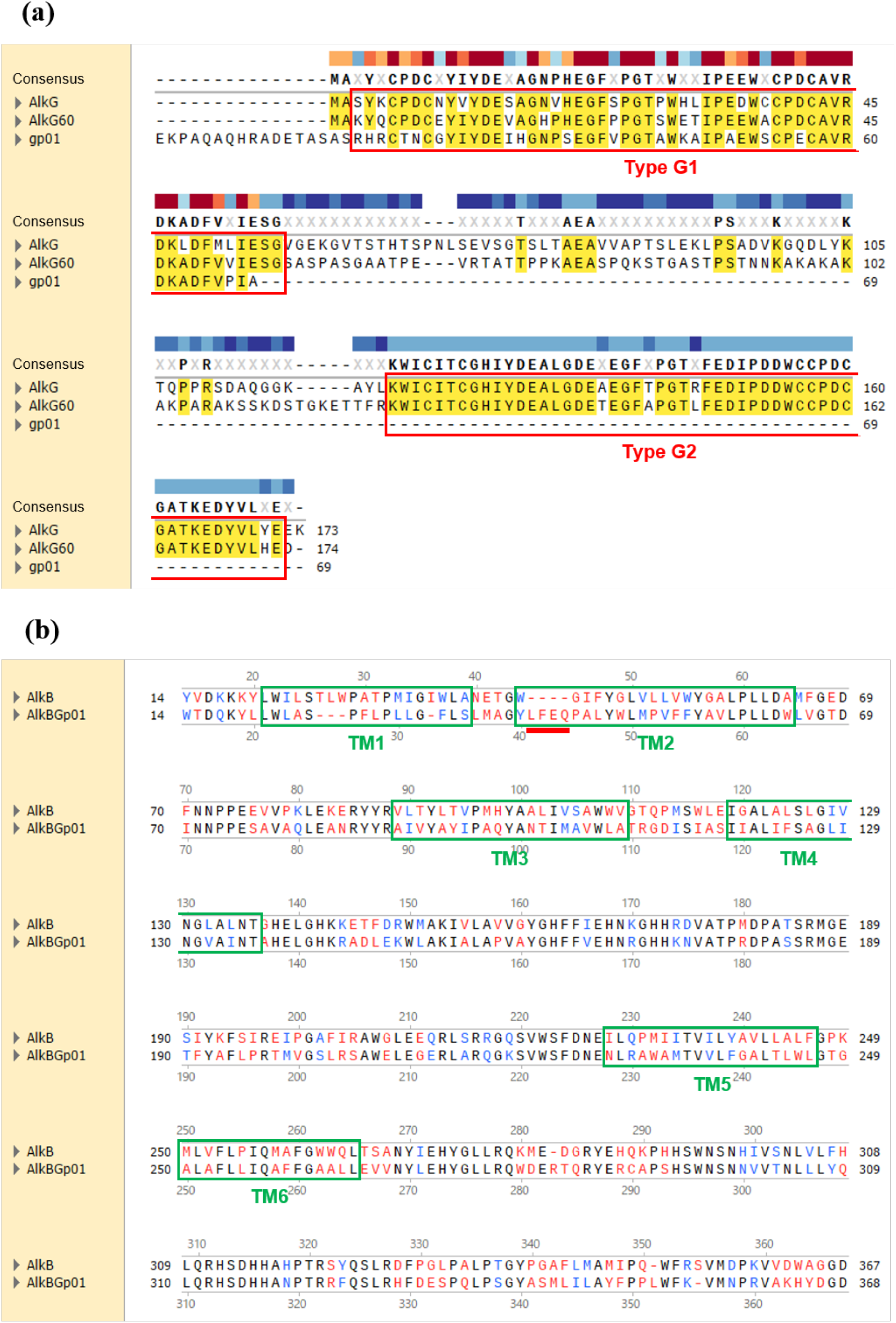
Sequence alignments of AlkG, AlkG60, and the C-terminal rubredoxin domain of AlkBGp01 (a), and AlkB and AlkBGp01 (b).

